# Lop-eared rabbits have more aural and dental problems than erect-eared rabbits: a rescue population study

**DOI:** 10.1101/671859

**Authors:** Jade C. Johnson, Charlotte C. Burn

**Affiliations:** The Royal Veterinary College, Hawkshead Lane, North Mymms, AL9 7TA, UK

**Keywords:** Animal welfare, Conformation, Dental pathology, Otitis, Rabbits, Veterinary Research

## Abstract

This research aimed to assess whether rabbits having lop-ears is a welfare issue by investigating the occurrence of aural and dental pathology in lop-eared compared with erect-eared rabbits.

Thirty rabbits (15 lop-eared and 15 erect-eared) from a rabbit-only rescue shelter were examined. An otoscope was used to visualise the ear canals and mouth. Samples were taken from each ear to examine for mites, bacteria and yeast. Medical records were also examined.

Lop-eared rabbits showed statistically significantly more frequent ear canal stenosis, higher scores of cerumen and erythema, and more frequent potential pain response during ear examination, compared with erect-eared rabbits. We also found statistically significantly more frequent incisor pathology, molar overgrowth, molar sharpness, molar spurs and history of veterinary dental treatment in lop-eared compared with erect-eared rabbits. The effect sizes were often large. Age was not statistically significant between the lop- and erect-eared rabbit groups.

Thus, lop-eared rabbits are at an increased risk of aural and dental pathology. This brings into debate the ethics of breeding and buying lop-eared rabbits, as they are more likely to suffer conditions that negatively impact welfare, such as pain, and potentially deafness and difficulty eating.

## Introduction

Breeding animals for extreme traits is coming under scrutiny, and has thus far mostly focused on dogs^1–3^. However, the ethics of breeding extreme rabbit conformations, such as pendulous ears and brachycephaly, is also starting to be debated^4, 5^. Lops are among the most common pet rabbit breed groups, representing 57% (n=58/102) and 36% (n=449/1254) of pet rabbits in England according to two surveys by Mullan and Main (2006) and Rooney et al. (2014), respectively. They also recently comprised 58.1% of known rabbit breeds advertised via Pets4Homes, a popular free pet advertisement and information website in the UK^6^. Thus, if the lop-eared phenotype is associated with suffering, it would constitute a prevalent rabbit welfare issue. However, clear evidence of whether this is the case is lacking.

Domestication of the wild rabbit has created at least nine breeds with a lop-eared phenotype, as it is a heritable trait^7, 8^. The wild type has erect and mobile pinna due to three interdigitating auricular cartilages that provide a scaffold for the vertical ear canal^9, 10^. In contrast, lop-eared breeds have 3-5mm of soft tissue between the proximal acoustic meatus cartilage ring and the tragal cartilage of the distal ear canal, causing the pinna to fold and become pendulous^10, 11^.

Charles Darwin first noted how artificial selection for pendulous pinnae in rabbits was associated with altered skull morphology^12^. Cordero and Berns (2016) found supportive evidence of this in natural woodrat populations that have likely undergone selection for long ears, finding substantial covariance of ear length and shape of the neighbouring auditory meatus^13^. This means that the lop-eared phenotype could not just affect aural function directly, but also other aspects of cranial health, such as dental function.

### Potential consequences for aural health

Lop-eared rabbits have increased stenosis (narrowing) of their ear canals^9, 14, 15^. This, along with ear canal occlusion by the pinna, is believed by some veterinary professionals to reduce airflow and hinder expulsion of cerumen (or ‘earwax’)^10, 11, 14^. Cerumen accumulation at the base of the ear in lop-eared rabbits has been suggested as a predisposing factor for otitis externa, caused by bacterial or yeast overgrowths^11, 14, 16, 17^. Mite infestation with *Psoroptes cuniculi* can also cause otitis externa, usually with extensive crusting of the pinna^10^. Veterinary professionals have observed that otitis externa can cause head shaking, ear scratching, pain, depression and inappetance in rabbits^18, 19^, and if it progresses to otitis media and interna then head tilts and neurological deficits can occur^19^. Veterinary opinion suggests that deafness can result from excess cerumen accumulation, otitis interna, or chronic otitis externa or media, at least in cats and dogs^20^, so it is possible that this could occur in rabbits also. Additionally, if the length of the lop ear is great, such as in English lops, then there could be increased risk of trauma to the pinnae^5^.

Despite the above veterinary and animal welfare expertise suggesting impacts of the lop-eared morphology on aural health, there has been limited empirical research into the issue; we found just one empirical study specifically comparing lop-eared and erect-eared rabbits, which solely investigated aural Malassezia yeast variation, and found no significant difference^21^. Thus, there is currently little evidence to argue the ethics of breeding and buying lop-eared rabbits. Veterinary opinion is mostly formed on the basis of experience with populations requiring veterinary attention, so it is unknown whether the same associations exist in otherwise healthy rabbits. However, it is possible that they do, because canine research has found that dogs with pendulous ears have a higher risk of otitis externa and an increased risk of *Malassezia species* infection, compared with dogs with erect-ears^22–24^. Thus, we hypothesise that these findings may also extend to rabbits with lop-ears.

### Potential consequences for dental health

Veterinary professionals have suggested that the altered skull morphology of lop-eared rabbits can cause congenital malocclusion of the dental arcades, such as maxillary brachygnathism (relative shortening of the lower jaw)^25^. This is also believed to occur in brachycephalic and dwarf breeds including the Netherland dwarf^25, 26^. Rabbit teeth grow continuously, so reduced attrition caused by malocclusion (i.e. if the upper and lower teeth are mis-aligned) can cause the teeth to overgrow^10, 27^. This has animal welfare implications because, if the teeth are not worn down appropriately then incisor overgrowth, elongated molar tooth roots and molar spurs can occur^27^. Rabbit incisors can grow at 2-2.4mm per week so problems can rapidly occur and reoccur after treatment, especially if the problem is due to malocclusion and thus cannot be rectified by husbandry changes^28^. Maxillary root overgrowth can penetrate the nasolacrimal duct and present as epiphora (watering of the eye) and secondary dacrocystitis (infection of the lacrimal sac)^28–30^. Root elongation can put pressure on sensory nerves innervating the teeth causing pain^28^. Further pain can occur if overgrown incisors penetrate the soft tissue of the lips, and mandibular and maxillary molar spurs can lacerate the lingual and buccal mucosa respectively^28^.

Dental disease is a chronic condition and the resulting oral pain and physical restrictions of overgrown teeth can lead to inappetance, poor body condition, and if severe can cause mortality through gut stasis^29, 31^. Other presentations of dental disease due to oral discomfort are hypersalivation, and a lack of grooming or caecotrophy that can lead to flystrike^27, 29, 31^.

### Aims and hypotheses

This research evaluated the number and type of aural and dental abnormalities in lop-eared versus erect-eared rabbit groups in a non-veterinary context: a rabbit-only rescue centre. We used two complementary sources of information: standardised direct observations of the rabbits, and medical records for the same rabbits; the latter were more heterogeneous and clinically driven than the observations, but could indicate past issues in the rabbits.

We hypothesised that, if the lop-eared phenotype leads to functional impairment, there would be significantly more aural pathology, such as ear canal stenosis, cerumen accumulation and inflammation, and dental pathology, such as incisor overgrowth and molar spurs, in lop-eared rabbits than in erect-eared rabbits.

## Materials and Methods

### Animals and husbandry

A convenience sample of fifteen lop-eared and fifteen erect-eared rabbits were selected for examination at a rabbit rescue centre. The sample size of 30 was chosen on the basis of feasibility, because the observations required completion within a relatively short time period, and without disrupting the husbandry routines of the rabbits at the shelter. The rabbits were selected by the rescue manager, who attempted to include a wide range of breeds and ages (where ages were known), and only included rabbits amenable to handling (for animal welfare, and health and safety reasons). She was not aware of the hypothesis, but she was aware that we would examine the ears and teeth of the rabbits as well as general health, and we did specify an equal number of lop- and erect-eared rabbits.

The rescue centre was capable of holding approximately 100 rabbits. All rabbits observed were housed in outdoor enclosures measuring at least 8ft × 6ft, which varied but typically comprised a large aviary with a hutch or shelter. Most rabbits (22/30) were housed in opposite sex neutered pairs, whilst eight were singly housed, and environmental enrichment included tunnels and chew toys. Water was provided *ad libitum* from bottles, and the diet consisted of a twice daily supply of hay, pellets (Burgess™, Pickering), and fresh leaves and vegetables.

Data were collected about each rabbit, including ear type (lop or erect), breed, face shape, sex, age, weight, and body condition score^32^. Face shape was assessed by subjectively observing the approximate length and breadth of the skull, and subsequently categorised into brachycephalic (broad, short skull, particularly seen in breeds such as the Netharland Dwarf and Lionhead), mesocephalic (moderately short and broad skull) and doliocephalic (narrow and long skull, as seen in the wild type) based on the independent opinion of two observers. Data were collected on standardised recording sheets (Supplementary material). The study received ethical approval from the Royal Veterinary College’s Clinical Research Ethical Review Board (URN 2015 1372).

### Aural health examination

Firstly, an observer (JCJ) stood quietly, approximately 1m from the enclosure, for 5 min to record presence or absence of head shaking and ear scratching, and the number of bouts of these. Then, the rabbit was gently restrained by an assistant whilst the observer performed a subjective clinical examination as summarised in Table 1 (details can be seen in supplementary information). Unfortunately, it was not possible to test inter-observer reliability during the examinations, as the available assistants were not medically trained so could not assess the ears or teeth. During direct observations, the observer could also not be blind to ear shape, so to minimise bias, this was explicitly discussed before observations started; we examined the possibility that lop-eared rabbits having conformational problems could be erroneous, especially since the only scientific comparison to date had found no cytological issues^21^, and that science was important for debunking incorrect perceptions. We were also aware of methods for interpreting and publishing studies where no statistical significance was founde.g. ^33, 34^, which minimised any incentive to find differences where none may exist.

**Table 1.**
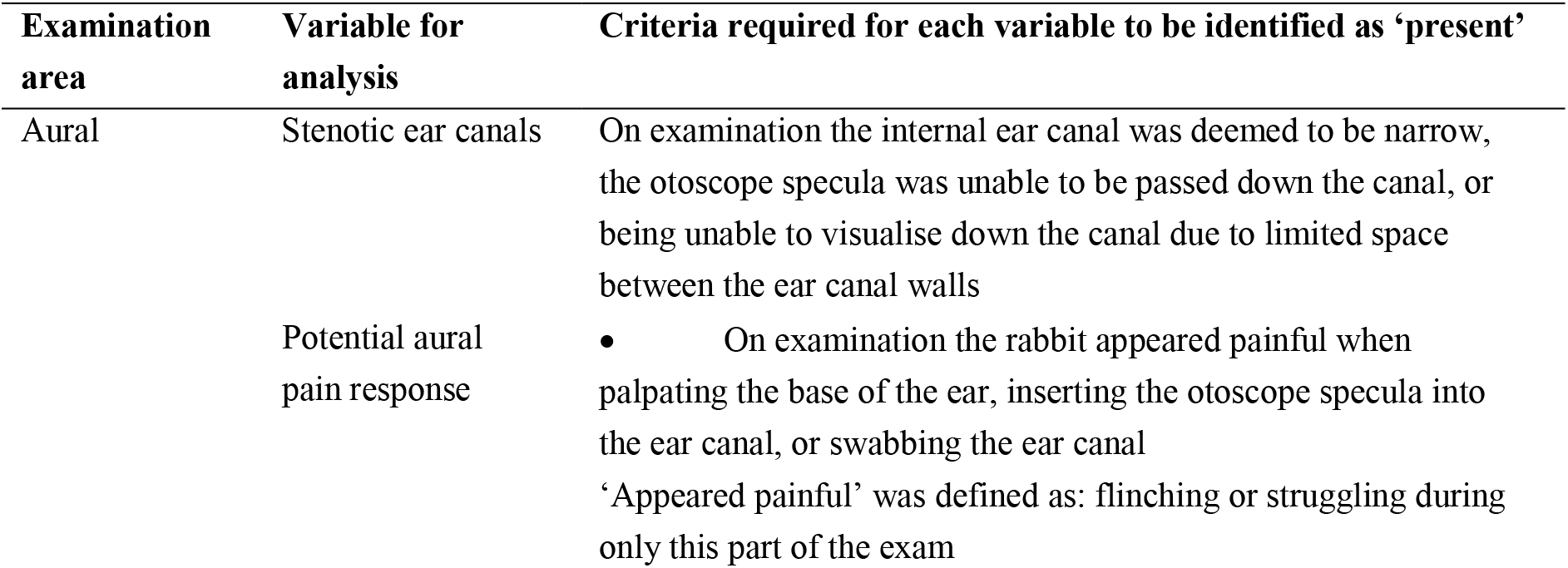
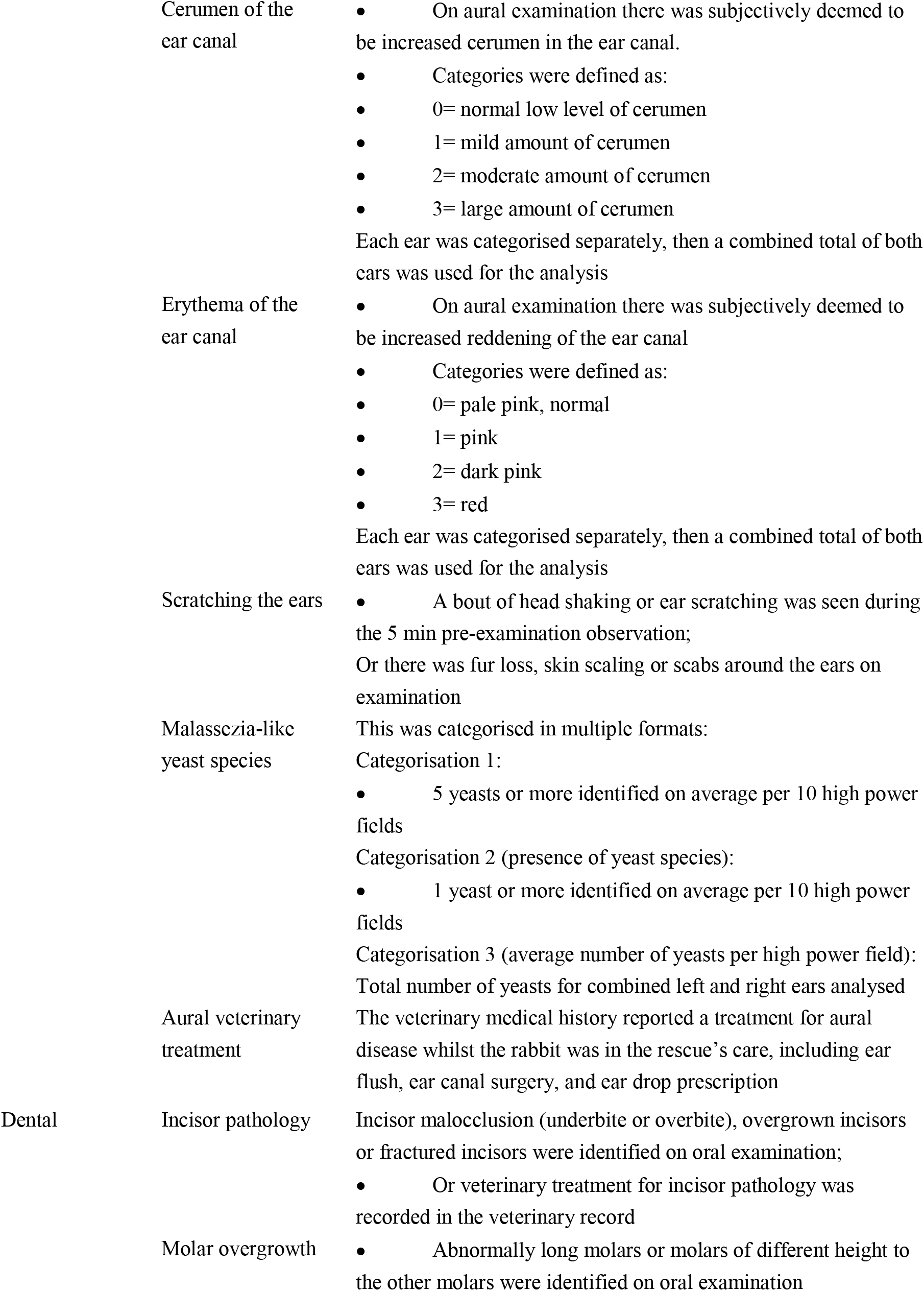
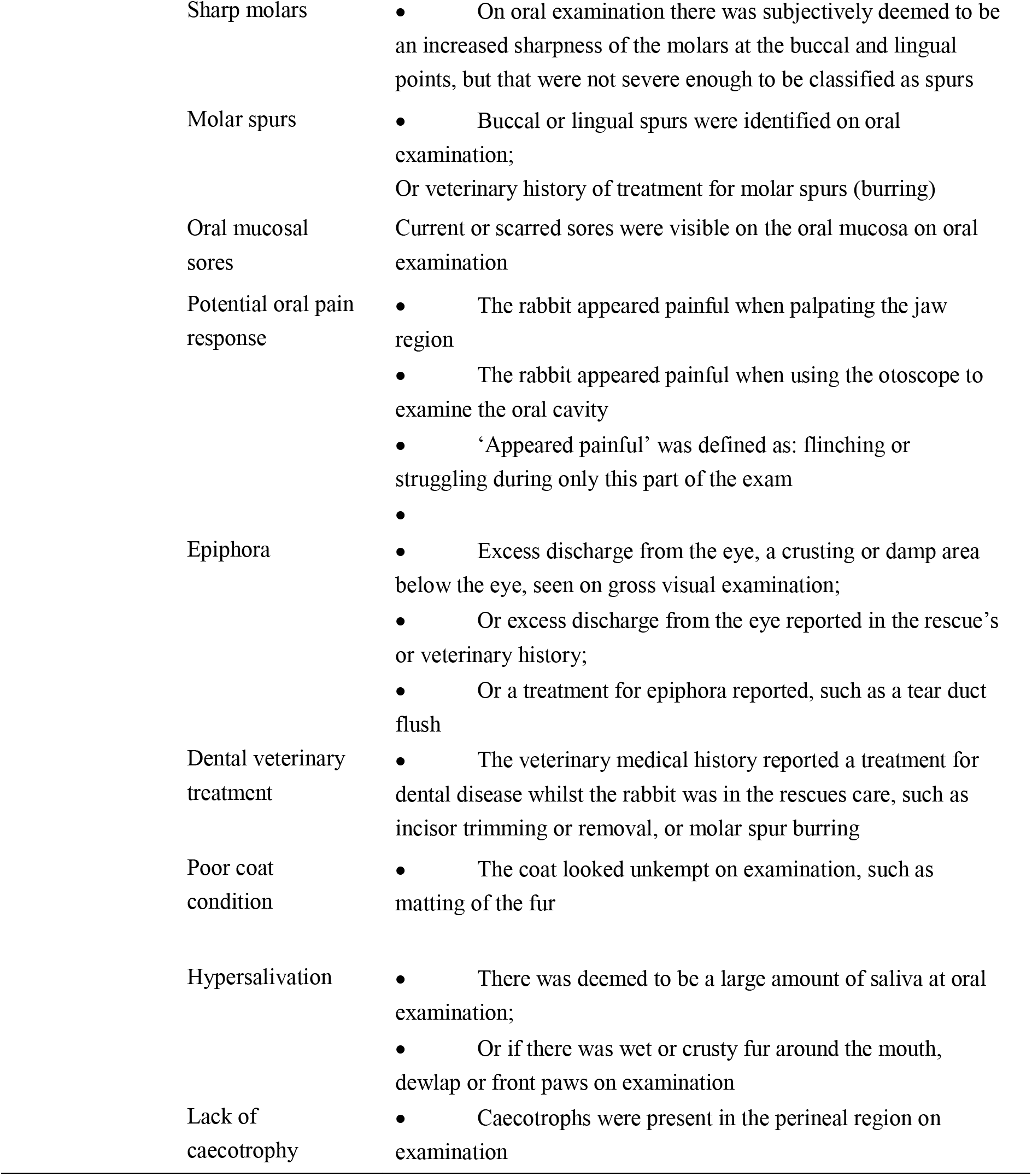
The descriptive criteria for each variable investigated statistically. The criteria are arranged such that aural pathology is first described, followed by dental pathology. Full details of the data collected are provided in the Supplementary information.

| Examination area | Variable for analysis | Criteria required for each variable to be identified as 'present' |
| --- | --- | --- |
| Aural | Stenotic ear canals | On examination the internal ear canal was deemed to be narrow, the otoscope specula was unable to be passed down the canal, or being unable to visualise down the canal due to limited space between the ear canal walls |
|  | Potential aural pain response | On examination the rabbit appeared painful when palpating the base of the ear, inserting the otoscope specula into the ear canal, or swabbing the ear canal<br>'Appeared painful' was defined as: flinching or struggling during only this part of the exam |
|  | Cerumen of the ear canal | On aural examination there was subjectively deemed to be increased cerumen in the ear canal.<br>Categories were defined as:<br>0= normal low level of cerumen<br>1= mild amount of cerumen<br>2= moderate amount of cerumen<br>3= large amount of cerumen<br>Each ear was categorised separately, then a combined total of both ears was used for the analysis |
|  | Erythema of the ear canal | On aural examination there was subjectively deemed to be increased reddening of the ear canal<br>Categories were defined as:<br>0= pale pink, normal<br>1= pink<br>2= dark pink<br>3= red<br>Each ear was categorised separately, then a combined total of both ears was used for the analysis |
|  | Scratching the ears | A bout of head shaking or ear scratching was seen during the 5 min pre-examination observation;<br>Or there was fur loss, skin scaling or scabs around the ears on examination |
|  | Malassezia-like yeast species | This was categorised in multiple formats:<br>Categorisation 1:<br>5 yeasts or more identified on average per 10 high power fields<br>Categorisation 2 (presence of yeast species):<br>1 yeast or more identified on average per 10 high power fields<br>Categorisation 3 (average number of yeasts per high power field):<br>Total number of yeasts for combined left and right ears analysed |
|  | Aural veterinary treatment | The veterinary medical history reported a treatment for aural disease whilst the rabbit was in the rescue's care, including ear flush, ear canal surgery, and ear drop prescription |
| Dental | Incisor pathology | Incisor malocclusion (underbite or overbite), overgrown incisors or fractured incisors were identified on oral examination;<br>Or veterinary treatment for incisor pathology was recorded in the veterinary record |
|  | Molar overgrowth | Abnormally long molars or molars of different height to the other molars were identified on oral examination |

|  |  |
| --- | --- |
| Sharp molars | On oral examination there was subjectively deemed to be an increased sharpness of the molars at the buccal and lingual points, but that were not severe enough to be classified as spurs |
| Molar spurs | Buccal or lingual spurs were identified on oral examination;<br>Or veterinary history of treatment for molar spurs (burring) |
| Oral mucosal sores | Current or scarred sores were visible on the oral mucosa on oral examination |
| Potential oral pain response | The rabbit appeared painful when palpating the jaw region<br>The rabbit appeared painful when using the otoscope to examine the oral cavity<br>‘Appeared painful’ was defined as: flinching or struggling during only this part of the exam |
| Epiphora | Excess discharge from the eye, a crusting or damp area below the eye, seen on gross visual examination;<br>Or excess discharge from the eye reported in the rescue’s or veterinary history;<br>Or a treatment for epiphora reported, such as a tear duct flush |
| Dental veterinary treatment | The veterinary medical history reported a treatment for dental disease whilst the rabbit was in the rescues care, such as incisor trimming or removal, or molar spur burring |
| Poor coat condition | The coat looked unkempt on examination, such as matting of the fur |
| Hypersalivation | There was deemed to be a large amount of saliva at oral examination;<br>Or if there was wet or crusty fur around the mouth, dewlap or front paws on examination |
| Lack of caecotrophy | Caecotrophs were present in the perineal region on examination |

The bases of the ears were palpated for swelling and the skin checked for evidence of scratching. The external pinnae were observed for crusting and inflammation (erythema, heat and swelling). An otoscope was used to visualise the ear canal to look for erythema, stenosis, increased cerumen, abnormal masses and potential pain responses such as flinching or struggling during only this part of the exam: the examination was shortened and not forced if this occurred, for animal welfare reasons (if the rabbit showed this response generally upon initiation of the exam then it was deemed to be due to fear, rather than pain, and the rabbit was excluded from the study due to it being unable to be examined). The otoscope heads were sterilised in a chlorhexidine solution between subjects.

Two samples were taken from each ear by gently inserting a sterile cotton swab into the external part of the ear canal and rotating fully three times. The samples were labelled with codes so that the observer could examine them blind to whether they originated from a rabbit with lop or erect ears. One sample from each ear was put onto a glass microscope slide with liquid paraffin and then covered with a cover slip. These slides were evaluated microscopically the same day using low magnification (10X) and high contrast to assess for mites. The other sample was rolled onto a microscope slide, allowed to dry for 10 min and then stained using Diff-quik (Rapi-Diff II stain, Atom Scientific, Cheshire) as per the manufacturer guidelines. Cytological assessment began with 40X magnification to find a representative area of interest, which was then evaluated using oil immersion and 100X magnification, to count the average number of bacteria (cocci and rods), and yeasts across 10 microscopic high-power fields. Cocci were identified as well-defined, circular and blue stained, and rods were identified as well-defined, rod shaped and blue stained.

### Dental health examination

After conducting the aural examination, the rabbit’s face was palpated externally for facial swellings, bony protrusions along the mandible and maxilla, ability to close the mouth and symmetry of the face. The examiner gently separated the lips manually using fingers and the eight incisors were evaluated for pathology including malocclusion, overgrowth, fractures or abnormal appearance such as horizontal ridging of the tooth surface. Then an otoscope was used in the mouth to assess the molars for overgrowth, sharpness, spurs, fractures and variation in tooth height. The lingual and buccal soft tissues were examined using the otoscope for sores and scars indicating healed sores. Evidence of potential pain was monitored for, such as flinching or struggling during only this part of the exam, and the examination was again shortened if this occurred. Secondary signs of dental disease were looked for including epiphora, poor coat condition, hypersalivation (also seen as wet or crusty fur around the mouth and throat), and caecotrophs around the perineum.

### Medical history records

The medical history of the rabbits whilst at the shelter was checked using hard copies of veterinary records and the rescue centre’s weekly health check record, especially focusing on dental and aural disease and any treatments for these. This was carried out after all rabbit examinations were completed, so that the examinations were not biased by knowledge of existing medical conditions.

### Statistical analysis

Where necessary, rare categories of data were collapsed together with similar categories to enable statistical analysis, with the final categories as described in Table 1. IBM SPSS statistics version 24 was used to carry out binary logistic regression on binary dependent variables. The predictors used in the model were ear type, sex, weight and face shape. Five rabbits had unknown ages (three with erect ears and two with lop ears), so, because age was not a statistically significant predictor in any of the initial models, and because mean age showed no significant difference between both groups (as analysed using binary logistic regression: P=0.492), it was removed from the final models, allowing data from all 30 rabbits to be included. The following outcomes were compared in lop- and erect-eared rabbit groups using binary logistic regression: incisor pathology, molar overgrowth, sharp molars, epiphora, lack of caecotrophy, stenosis of the ear canal, potential aural pain, and the presence of yeast.

However, no rabbits from the erect-eared group at all were affected for the following variables: molar spurs, potential oral pain response, oral mucosal sores, poor coat condition, hypersalivation, veterinary dental treatment, aural veterinary treatment, and scratching the ears. This meant there were fewer than five values in one group, so a Fisher’s exact statistical test was carried out using GraphPad Prism Version 7.

For outcomes that were recorded as continuous scores, Mann-Whitney U statistical tests were carried out using GraphPad Prism Version 7. This was the case for the average number of yeasts per high power field, cerumen in the ear canal and erythema in the ear canal.

## Results

### Demographics

Rabbit signalment is shown in Table 2. The known ages of the erect-eared rabbit group ranged from 5 months to 9 years, whilst those of the lop-eared rabbit group ranged from 8 months to 7 years 9 months. The mean age was not statistically different between the lop- and erect-eared groups (P=0.492). The rabbits’ body weights ranged from 1.4kg to 3.5kg in both groups. The erect-eared breeds mainly consisted of cross breeds (n=10), followed by Lionheads (n=3), whilst Dwarf lops (n=3) and Mini Lops (n=3) were the most common lop-eared rabbits.

**Table 2.**
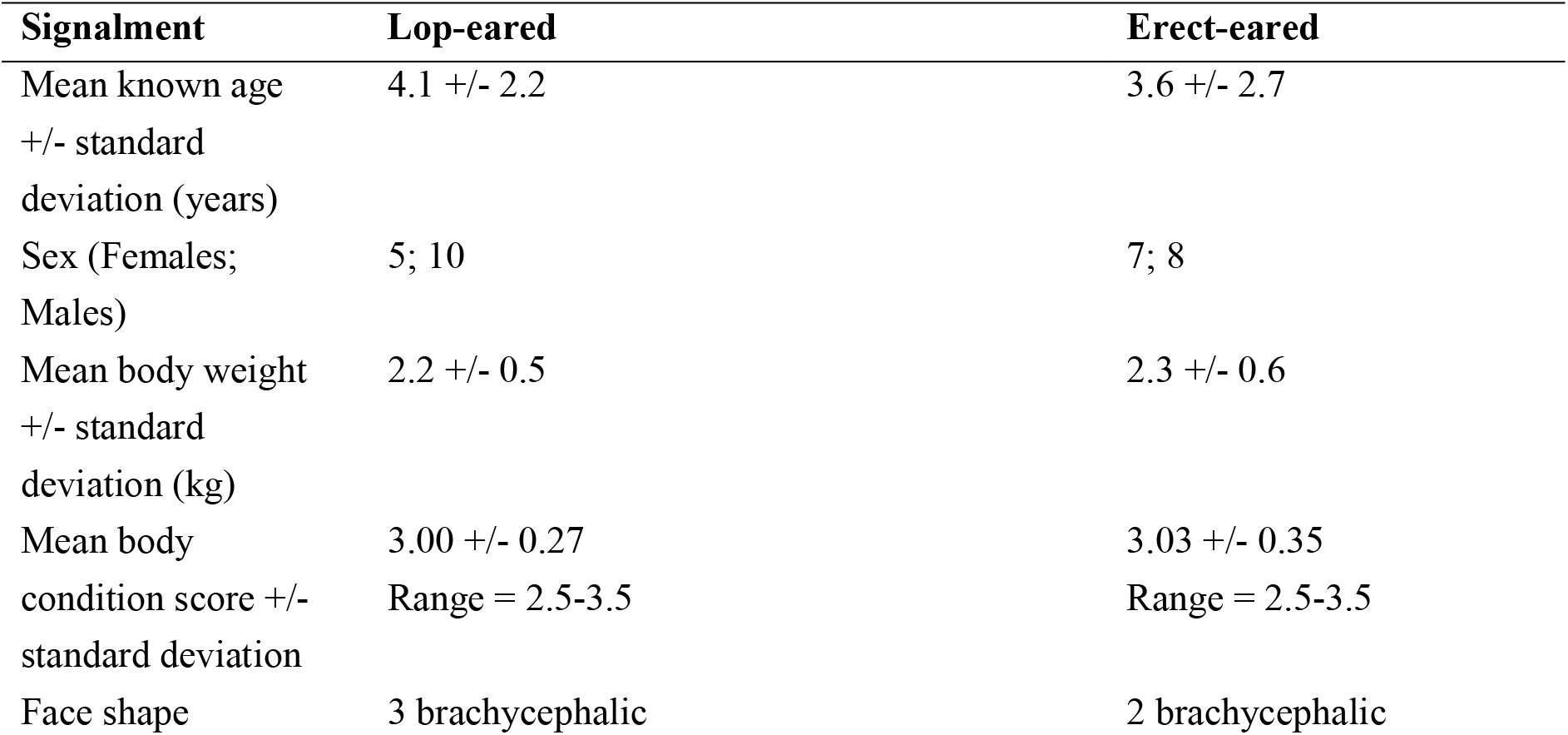
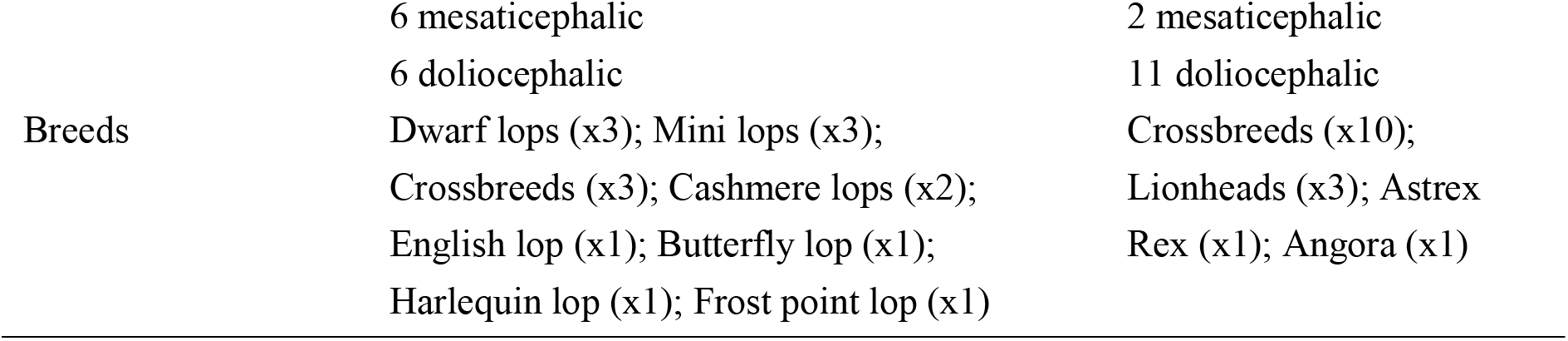
Demographic details of the lop-eared and erect-eared rabbit groups.

### Aural health results

Statistical results are summarised in Table 3. Lop-eared rabbits had approximately forty-three times higher odds of having stenotic ear canals (OR=42.7; 95% CI: 4.2-434; P=0.002; Figure 1) and a statistically significantly higher erythema score of the ear canal (U=50.5; N=15; P=0.004; Figure 2), compared with erect-eared rabbits. They also had a statistically significantly higher cerumen score in the ear canal compared with erect-eared rabbits (U=24; N=30; P<0.001; Figure 3).

**Table 3.**
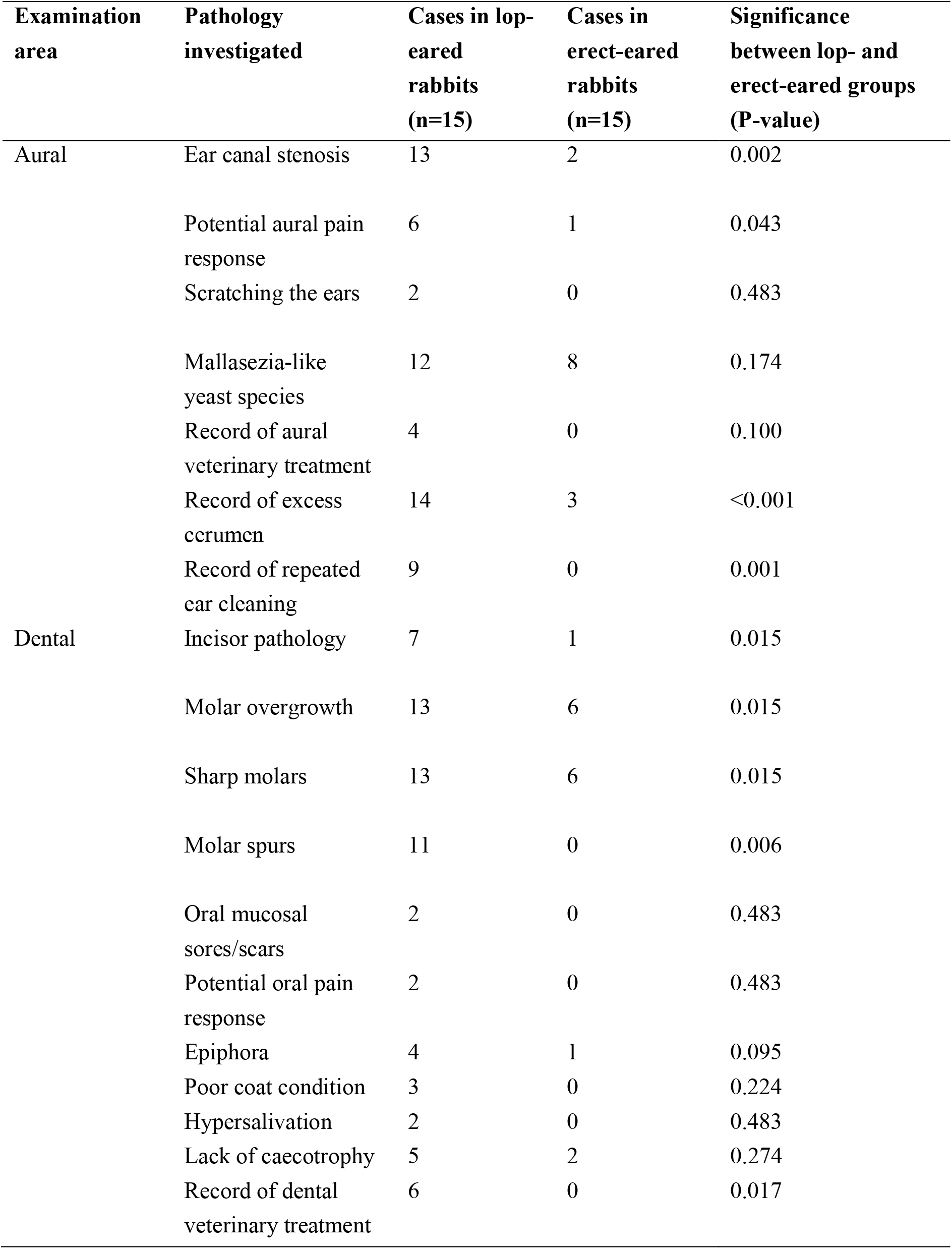
Prevalence of binary variables investigated in both lop and erect-eared rabbit groups and the significance of the difference between both groups.

**Figure 1.**
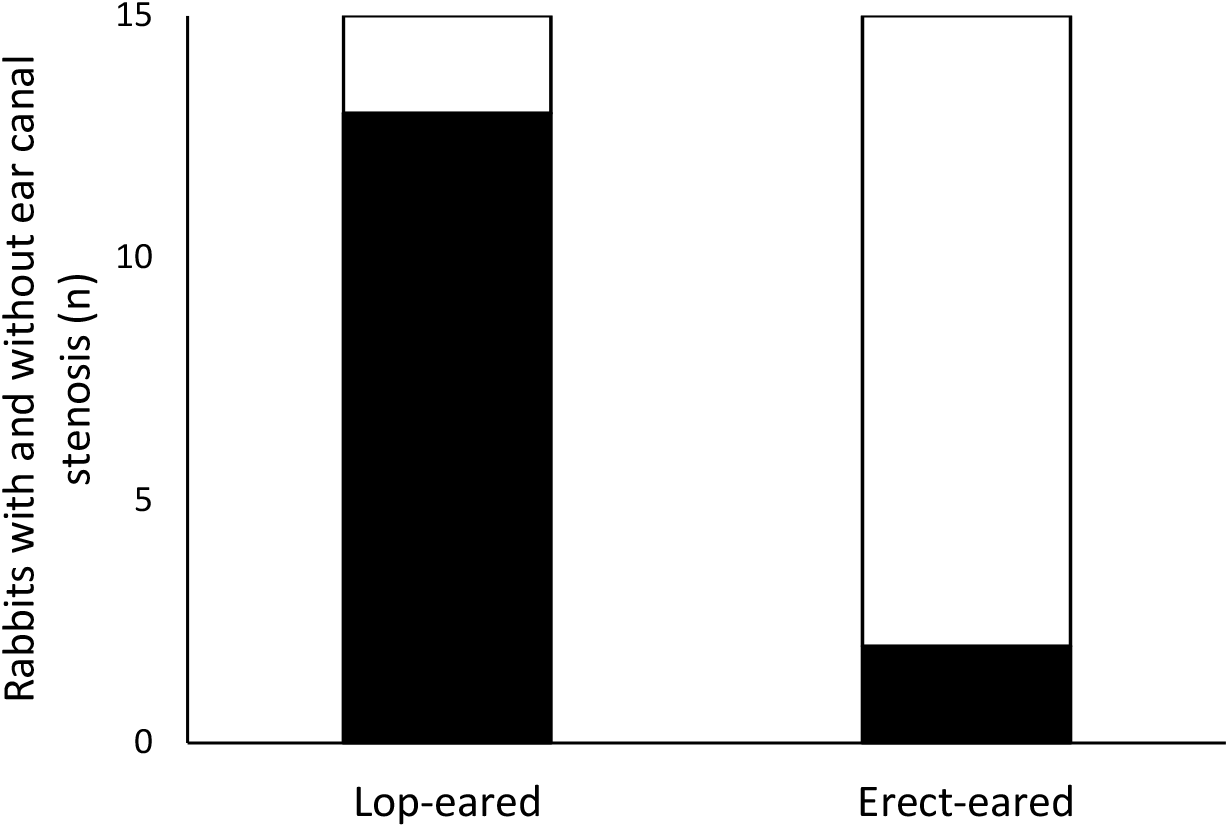
Number of lop- and erect-eared rabbits with and without ear canal stenosis. Black indicates presence of stenosis; white indicates absence of stenosis. Significantly more (13/15) lop-eared rabbits had ear canal stenosis compared with (2/15) erect-eared rabbits (P=0.002).

**Figure 2.**
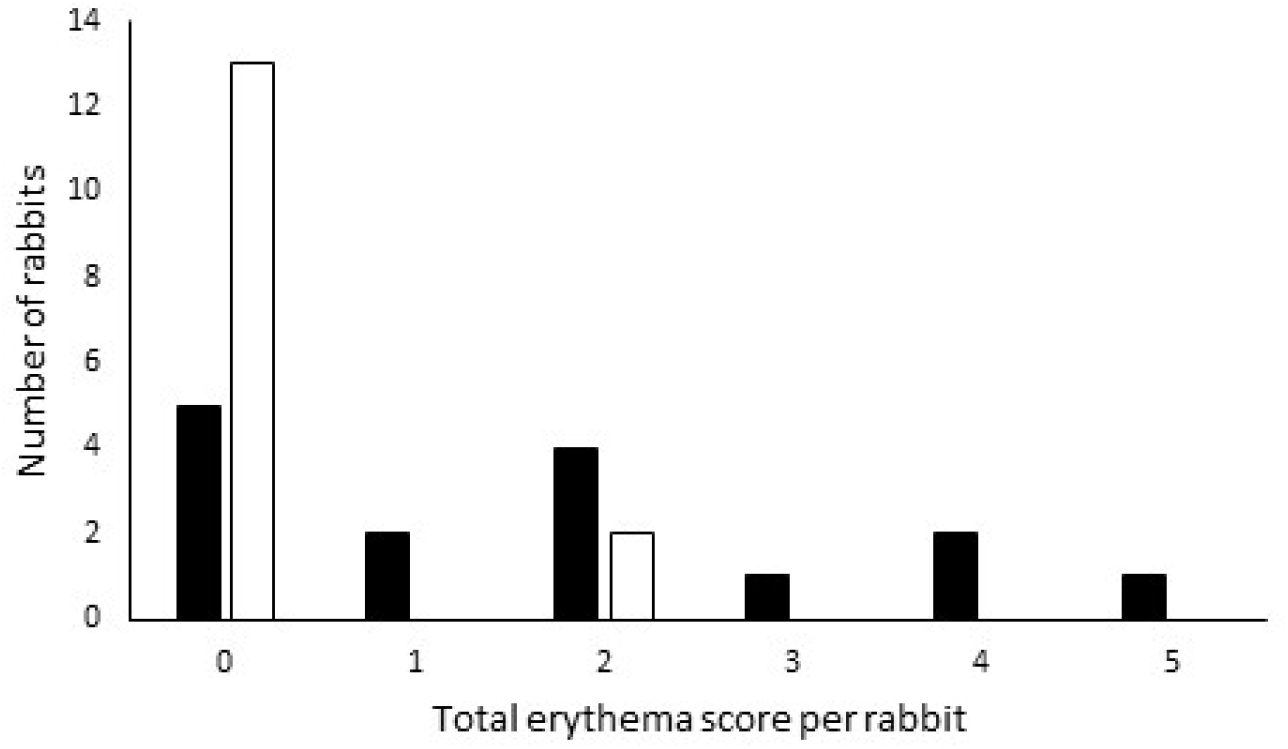
Frequency distribution of erythema scores in the ear canals of lop-eared and erect-eared rabbits. Black bars indicate lop-eared rabbits; white bars indicate erect-eared rabbits. Each ear was scored as follows: 0= pale pink, 1= pink, 2= dark pink, 3= red, and the added total for both ears per rabbit is shown.

**Figure 3.**
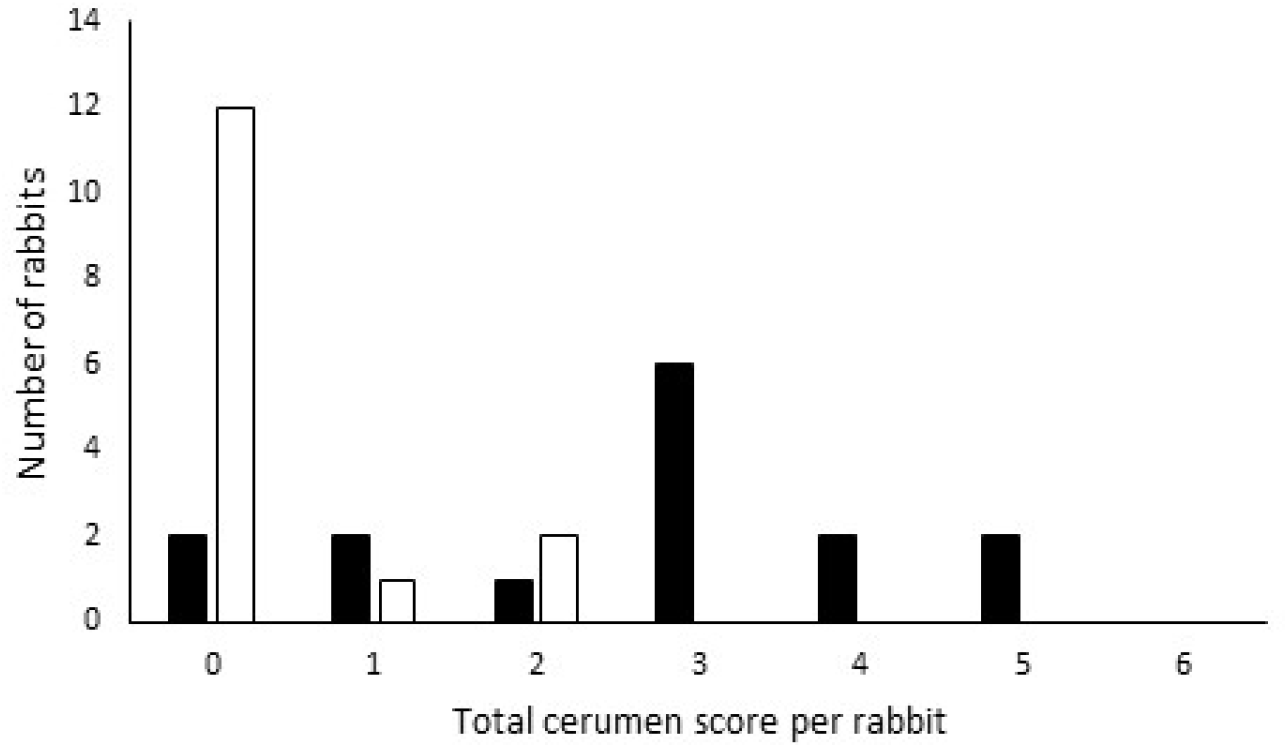
Frequency distribution of cerumen scores lop-eared and erect-eared rabbits. Black bars indicate lop-eared rabbits; white bars indicate erect-eared rabbits. Each ear was scored as follows: 0= normal low amount of cerumen, 1= mild amount of cerumen, 2= moderate amount of cerumen, 3= large amount of cerumen, and the added total for both ears per rabbit is shown.

Lop-eared rabbits had approximately fifteen times higher odds of demonstrating a potential pain response during ear examination than erect-eared rabbits (OR=14.8; 95% CI: 1.1-200.9; P=0.043; Figure 4).

**Figure 4.**
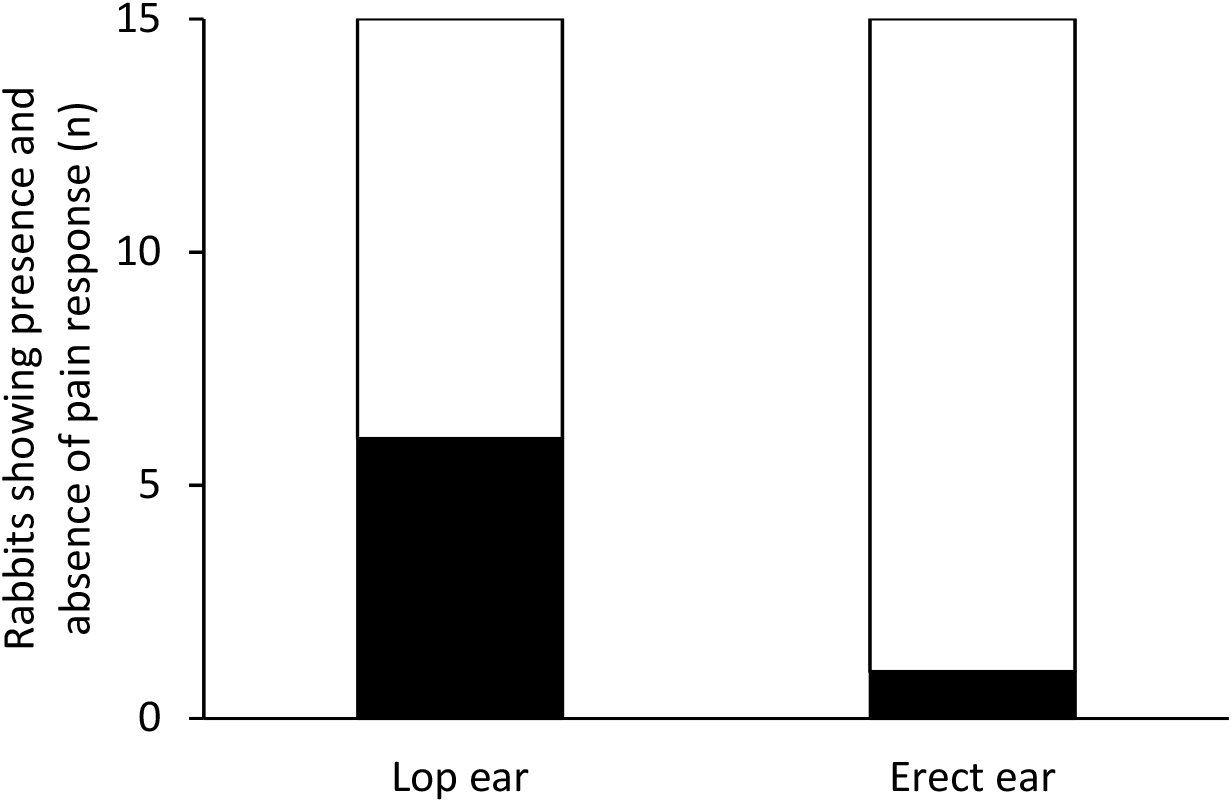
Frequency distribution of potential pain response during ear examination in lop- and erect-eared rabbits. Black indicated presence of a potential pain response during examination of at least one ear; white indicates absence of a pain response.

The health records revealed that 14/15 of the lop-eared rabbits had at least one recording of excess wax during health checks by the rescue staff, compared with 3/15 of the erect-eared rabbits (Fisher’s Exact X^2^= 13.575; DF = 1; P < 0.001). The rescue carried out repeated ear cleaning in 9/15 of the lop-eared rabbits, compared with none of the erect-eared rabbits (Fisher’s Exact X^2^= 10.159; DF = 1; P = 0.001).

No statistically significant difference was found between lop-eared and erect-eared rabbits for evidence of scratching (fur loss or scaling) around the ears (2/15 versus 0/15, respectively), or ear pathology requiring treatment whilst at the rescue centre (4/15 versus 0/15, respectively). None of the rabbits had crusting of the external pinna, masses in the ear, head shaking or scratching in pre-examination observation or a head tilt. No statistical significance was found for any of the other predictors used in the binary logistic regression (face shape, sex or weight) for any of the aural pathology outcomes.

Microscopy revealed the presence of yeast species in some rabbits (12/15 lop-eared rabbits versus 8/15 erect-eared rabbits), which were identified as *Malassezia cuniculi* based on morphology described in other research^21, 24, 35^. The presence/absence, and also number of, malassezia-like yeasts found on microscopy were not statistically significantly different between lop- and erect-eared rabbits.

Only one mite was found, identified as Psoroptes cuniculi, in the ear of an erect-eared rabbit. This rabbit had no associated clinical signs or previous history of aural problems. No rod bacteria were identified, but cocci were found in three lop-eared rabbits and two erect-eared rabbits.

### Dental health results

Lop-eared rabbits had approximately twenty-three times higher odds of incisor pathology compared with erect-eared rabbits (OR=23.3; 95% CI: 1.9–293.2; P=0.015). They also had approximately twelve times higher odds of having molar overgrowth (OR=12; 95% CI: 1.6-88.9; P=0.015), thirteen times higher odds of molar sharpness (OR=13.4; 95% CI: 1.7-107.2; P=0.015) and were statistically significantly more likely to have molar spurs (attributable risk=0.4667; P=0.006), compared with erect-eared rabbits (Table 3; Figure 5).

**Figure 5.**
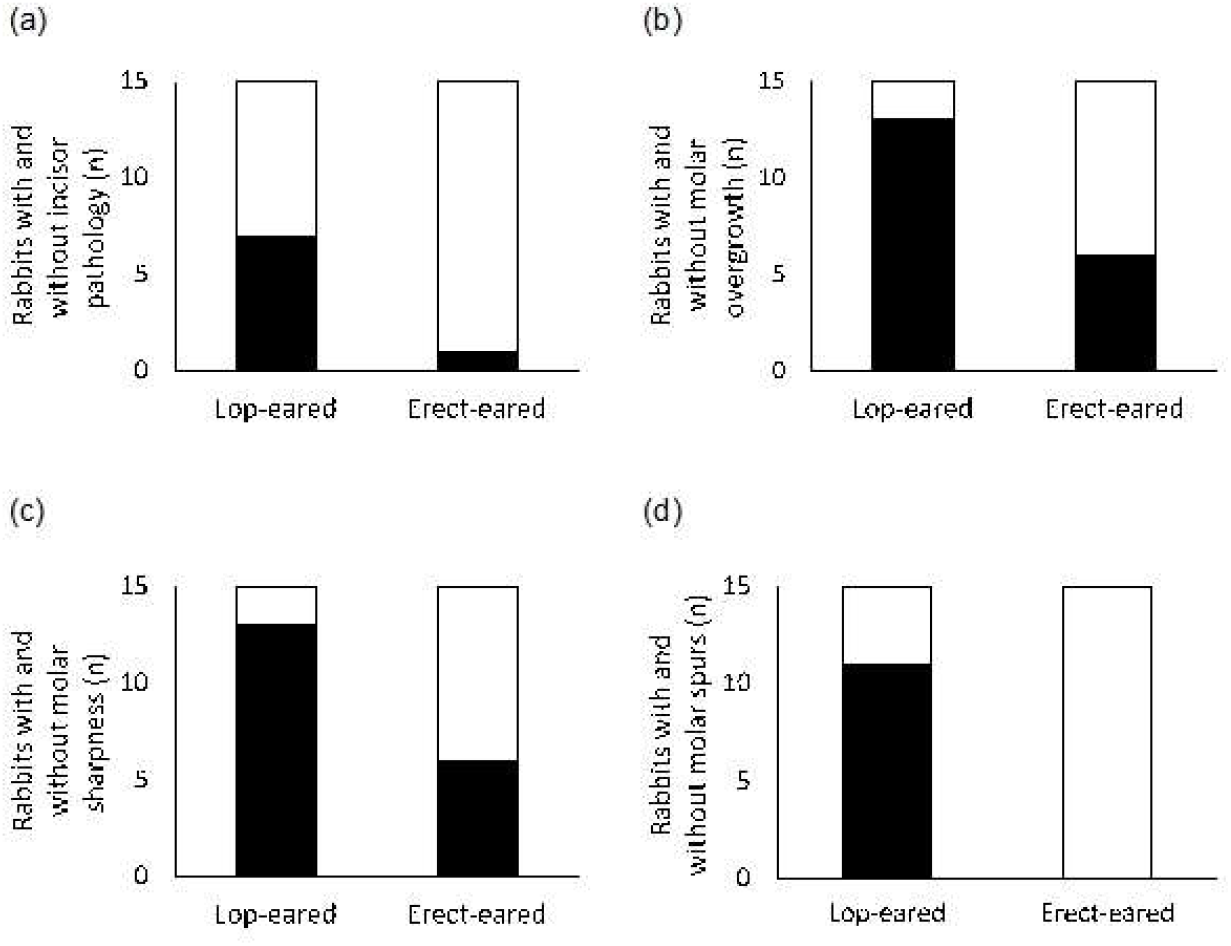
Frequency distribution of dental pathology in lop- and erect-eared rabbits. Black indicates presence of pathology; white indicates its absence. Figure (a) shows incisor pathology, (b) molar overgrowth, (c) molar sharpness, and (d) molar spurs.

Veterinary records showed that 8/15 of the lop-eared rabbits had tooth abnormalities, compared with none of the erect-eared rabbits (attributable risk= 0.53, 95% CI: 0.17-0.78; P=0.002). Additionally, 6/15 of the lop ears had been for a dental whilst at the rescue, whereas none of the erect-eared rabbits had, which was again statistically significant (attributable risk=0.4; 95% CI: 0.06-0.67; P=0.017).

On the other hand, statistically significant differences between lop and erect-eared rabbits were not found for oral mucosal sores, potential oral pain response, epiphora, poor coat condition, hypersalivation, or lack of caecotrophy, although the trends were for all of these to be slightly more likely in lop-eared rabbits (Table 3). No rabbits had an inability to close the mouth or fractured teeth.

No statistical significance was found for any of the other predictors used in the binary logistic regression (face shape, sex or weight) for any of the dental pathology outcomes.

## Discussion

The results confirmed that lop-eared rabbits did indeed have a statistically significantly higher level of aural pathology, including stenotic ear canals, potential pain response during aural examination, increased levels of cerumen and erythema of the ear canal. Similarly, there was also a statistically significantly higher level of dental pathology, including incisor pathology, molar overgrowth, molar sharpness, molar spurs and veterinary dental treatment, in lop-eared compared with erect-eared rabbits. Both the direct observations and the medical records showed that the lop-eared rabbits had statistically significantly more aural and dental pathologies than did erect-eared rabbits.

The use of a rescue population may of course not represent the general population of pet rabbits. It is difficult to know if rescue rabbits would be affected more than pet rabbits by the issues investigated here, or less. On one hand, rabbits in the rescue centre may be more prone to health problems if they were given to the rescue because their owners could not afford veterinary bills, or due to neglect and being fed an inappropriate diet. On the other hand, the rescue staff were highly knowledgeable about rabbit health and carried out weekly health checks, so current severe dental or aural disease was not found during the clinical examination of this study. In either case, lop- and erect-eared rabbits would have been affected by these factors to a similar extent. Thus, the finding that lop-eared rabbits were prone to the aural and dental pathology investigated here constitutes a welfare concern associated with breeding and buying lop-eared rabbits, for reasons described below in more detail.

### Aural pathology

The increased erythema, ceruminous discharge and stenosis of the ear canal found in the lop-eared rabbits here, could indicate a higher frequency of otitis externa. This would concur with two canine studies that found higher prevalences of otitis externa in dogs with pendulous ears^22, 23^. In many of the rabbits, this appeared to be a chronic condition as the medical records showed that 9/15 of the lop-eared rabbits required repeated ear cleaning whilst at the rescue.

Thus, the welfare consequences of a rabbit having lop-ears include pain, as indicated by statistically significantly increased pain responses during examination of lop-ears. Additionally the higher frequency of signs consistent with otitis externa found in the lop-eared compared with the erect-eared rabbits, suggest potential for pain, auditory deficit or even deafness^18, 19^, which in turn increases the vulnerability of the animal to threats and could cause anxiety^20^. Deafness itself could not be tested in the current study, but this could be assessed in future studies using Transient Evoked Otoacoustic Emissions testing, which has been successfully used as a less invasive and relatively inexpensive alternative to brainstem auditory evoked responses in puppies^36^. The rescue centre staff anecdotally believed that more of the lop-eared rabbits had auditory impairment than the erect-eared ones had, and unpublished research from our laboratory suggested that lop-eared rabbits showed more signs of anxiety in a novel object test than erect-eared rabbits^37^.

The results here suggest that the aetiology of any potentially associated auditory deficits would be multifactorial as it is in dogs and cats^20, 38^, with the over-hanging immobile pinnae, the stenotic ear canals, and the accumulation of cerumen all comprising physical barriers to sound perception. The middle and inner structures themselves may also be more prone to issues, such as tympanic membrane rupture and sensorineural damage, if repeated infection occurs.

The lack of a statistically significant difference in the presence of *Malassezia cuniculi* between lop and erect-eared rabbits could of course be due to the fairly small sample size here, but it agrees with the findings of Quinton et al. (2014), who reported no statistically significant difference due to ear type in 146 clinically healthy domestic pet rabbits. This could be explained by *Malassezia* being a normal coloniser of the rabbit ear canal, as in other species such as dogs and cats^17, 21^. In future, culture could offer a more sensitive method for quantifying *Malassezia* colonisation than cytology^39^. Campbell et al.^17^ found a positive correlation between the amount of ceruminous discharge and the culture of *Malassezia*. However, although the present study found higher levels of cerumen in lop-eared rabbits, the lack of difference in *Malassezia* colonisation suggests that this increased cerumen was due to difficulties in expulsion, potentially due to the anatomy of the lop-ear, rather than a *Malassezia* overgrowth^10^.

### Dental pathology

The increased presence of dental pathology found in lop-eared rabbits in this study partially supports results from Mullan and Main^40^, who found that dwarf lops were reported by their owners to have statistically significantly more dental abnormalities compared with other breeds (including some other lops), but only when diet and age were excluded from the model. However, the present results disagree with a Finnish study^41^ of 167 pet rabbits, which found no associations between lop-eared breeds and dental disease, although they did find statistically significantly more dental pathology in Lionhead rabbits, which are erect-eared, but brachycephalic. A possible explanation for the disagreement is that the study by Mäkitaipale et al.^41^ was carried out on healthy pet rabbits whose owners voluntarily signed up for the study, so those with known current dental problems may have been excluded from that study.

The rabbits in our study represent a convenience sample of a rescue population, but if our results can be replicated more widely, the increased risk of dental pathology for lop-eared rabbit welfare is concerning for several reasons. Rabbit dental pathology can lead to lesions of the mouth, pain, difficulty chewing food e.g. avoidance of, or prolonged chewing of, hard foods as seen in bears with dental pathology^42^, and possibly gastrointestinal problems following inadequately chewed food as found in humans^43, 44^, among other issues described earlier. The rescue staff caring for rabbits in the current study were more knowledgeable of and attentive to rabbit health issues than many owners, and health records indicated that they had already identified more dental issues requiring veterinary attention in lop-eared than erect-eared rabbits, before our study started. Indeed, the rabbits at the rescue centre underwent weekly health checks by experienced rescue staff with the use of an otoscope. Despite this enhanced care and monitoring of dental pathology, we observed statistically significantly more abnormalities in both incisors and molars in the lop-eared rabbits. The potentially associated secondary issues, such as trauma to the mouth and lips, hypersalivation, and pain responses, were too rare to reach statistical significance, but all showed trends towards being slightly more common in lop-eared rabbits.

The dental abnormalities here are unlikely to be due to diet, because there is currently no a priori reason to believe that lop-eared rabbits would have been fed poorer diets than erect-eared rabbits. Thus, they are likely to be caused at least partly by the skull and jaw conformations associated with the lop-eared phenotype. Future studies could help confirm this by observing random sample populations, and by utilising radiology to look at oral conformations, alongside oral examination if possible^29, 31, 41^.

### Conclusions

The results from this research support the hypothesis that lop-eared rabbits have more dental and aural pathology than erect-eared rabbits, which is concerning given the popularity of the lop-eared phenotype. This brings into debate the ethics of breeding and buying lop-eared rabbits, as they appear more likely to suffer from these conditions, which can be painful and often chronic or recurrent.

## Acknowledgements

The authors wish to thank the staff at The Rabbit Residence Rescue, Royston for allowing the inclusion of their rabbits in the project, Dr Anette Loeffler for her guidance on making and interpreting ear smears, Dr Anke Hendricks for her help with equipment and advice, Katie Lovell for supplying laboratory equipment, Katie Franklin for her assistance at the rescue and providing the list of rabbits, and Natalie Chancellor and Tomaso Carucci for their help handling the rabbits for examination. This research was carried out in part fulfilment of JCJ’s Bachelor of Veterinary Medicine with Intercalated BSc Bioveterinary Sciences degree, in year 5 of this course at the Royal Veterinary College. The Royal Veterinary College assigned the following ID number to this study: PPS_01851.

