## Supplemental Data Recording Sheet for "Lop-eared rabbits have more aural and dental problems than erect-eared rabbits: a rescue population study" (from another version of this preprint: version 1, 2019-06-14)

### Supplementary information: Rabbit examination recording sheet

Part 1: Rabbit details

1. Rabbit name: ……………………… Assigned number:……………………

2. Sex:………………

3. Age:………………

4. Lop eared □ Prick eared □ One lop ear, one prick ear □

5. Breed: …………………………

a. Dwarf □ Standard size □ Giant □

b. Weight:………..kg

6. Brachycephalic: Yes □ No □ Unsure □

7. Body condition score: …../5

8. Current diet: Pellets □ Cereal □ Hay □ Grass □ Fruit/veg □ Plants □ Knawing items (e.g wood) □ Soft mash □ Other:………………………………..

a. Is there a medical reason for the diet they are on?..............................................................

**Extra notes of interest about rabbit details**

……………………………………………………………………………………………

……………………………………………………………………………………………

……………………………………………………………………………………………

……………………………………………………………………………………………

……………………………………………………………………………………………

……………………………………………………………………………………………

……………………………………………………………………………………………

…………………………………………………………

Part 2: Aural exam

1. Normal: Yes □ No □ Unsure □
2. Gross exam of left ear:

a. Crusting of external pinna: Yes □ No □ Unsure □

i. If yes, how much of the pinna is covered in crusting?..........%

b. Inflammation (redness/heat/swelling) of external pinna: Yes □ No □ Unsure □

i. Severity: Lightly inflamed (pink, warm, light swelling) □ Intermediate inflammation (dark pink, warm-hot, medium swelling) □ Heavily inflamed (red, hot, large swelling) □ Other:………………………………………………….

c. Inflammation (redness/swelling) of internal ear canal: Yes □ No □ Unsure □

i. Severity: Lightly inflamed (pink, slight stenosis of ear canal) □ Intermediate inflammation (dark pink, obvious stenosis of ear canal but can still pass scope) □ Heavily inflamed (red, hot, complete stenosis of ear canal and cannot pass scope) □ Other:…………………………………………………………………….

d. Excess sebum: Yes □ No □ Unsure □

i. Severity: Mild increase in sebum □ Moderate increase in sebum □ Large increase in sebum □

ii. Location of excess sebum: Internal ear canal □ External ear canal opening/pinna □

e. Growths/masses in ear: Yes □ No □ Unsure □

i. How many masses are found in the ear?.................

ii. Description of mass/growth…………………………………………………………….

f. Evidence of potential pain (exaggerated shaking head & struggling away from scope): Yes  No  Unsure 

i. Unable to complete exam due to potential pain/refusal: Yes 

g. Other:……………………………………………………………………………………… ………………………………………………………………………………………………… …..

3. Gross exam of right ear:

a. Crusting of external pinna: Yes  No  Unsure 

i. If yes, how much of the pinna is covered in crusting?..........%

b. Inflammation (redness/heat/swelling) of external pinna: Yes  No  Unsure 

i. Severity: Lightly inflamed (pink, warm, light swelling)  Intermediate inflammation (dark pink, warm-hot, medium swelling)  Heavily inflamed (red, hot, large swelling)  Other:………………………………………………….

c. Inflammation (redness/swelling) of internal ear canal: Yes  No  Unsure 

i. Severity: Lightly inflamed (pink, slight stenosis of ear canal)  Intermediate inflammation (dark pink, obvious stenosis of ear canal but can still pass scope)  Heavily inflamed (dred, hot, complete stenosis of ear canal and cannot pass scope)  Other:…………………………………………………………………….

d. Excess sebum: Yes  No  Unsure 

i. Severity: Mild increase in sebum  Moderate increase in sebum  Large increase in sebum 

ii. Location of excess sebum: Internal ear canal  External ear canal opening/pinna 

e. Growths/masses in ear: Yes  No  Unsure 

i. How many masses are found in the ear?.................

ii. Description of mass/growth…………………………………………………………….

f. Evidence of potential pain (exaggerated shaking head & struggling away from scope): Yes  No  Unsure 

i. Unable to complete exam due to potential pain/refusal: Yes 

g. Other:………………………………………………………………………………………… ………………………………………………………………………………………… …………………………………….

4. Other signs of aural disease:

a. Head shaking/scratching on 5 min pre-exam observation: Yes  No 

i. If yes, how many bouts of head shaking/scratching were observed:……………

b. Head tilt: Yes  No 

i. In which direction: Left  Right 

c. Scratching of head/ears (marked by fur loss and scaling): Yes  No 

i. Left  Right  Both sides 

5. Smear results of left ear

a. Cytology on stained smear:

i. Bacterial infection: Cocci  Rods  Other:…………….. Unsure 

1. Number per hpf:………………

ii. Yeast seen: Malassezia  Other:…………….. Unsure 

1. Number per hpf:………………

iii. Other…………………………………………………………………………….

b. Plain smear

i. Mites (psoroptes cuniculi): Yes  No 

1. Number seen…………….

ii. Other:…………………………………………………………………………

6. Smear results of right ear

a. Cytology on stained smear:

i. Bacterial infection: Cocci ☐ Rods ☐ Other:…………….. Unsure ☐

1. Number per hpf:………………

ii. Yeast seen: Malassezia ☐ Other:…………….. Unsure ☐

1. Number per hpf:………………

iii. Other……………………………………………………………………………….

b. Plain smear

i. Mites (psoroptes cuniculi): Yes ☐ No ☐

1. Number seen…………….

ii. Other:………………………………………………………………………………

**Extra notes of interest on aural health**

………………………………………………………………………………………………

………………………………………………………………………………………………

………………………………………………………………………………………………

………………………………………………………………………………………………

………………………………………………………………………………………………

………………………………………………………………………………………………

………………………………………………………………………………………………

…………………………………………………………

Part 3: Dental exam

1. Normal: Yes □ No □ Unsure □

2. Swellings (e.g facial abscesses)/ bony lumps/tenderness along mandible/maxilla/facial asymmetry: Yes □ No □ Unsure □

a. If yes, state which & describe…………………………………………………………..……………………..

b. If yes, how many………………………..

c. If yes, location…………………………..

3. Normal degree of lateral movement of mandible: Yes □ No □ Unsure □

4. Inability to fully close the mouth: Yes □ No □ Unsure □

a. Is there a cause evident:………………………………………………………………………………………….

5. Incisor appearance:

a. Normal: Yes □ No □ Unsure □

b. Incisor ribbing: Yes □ No □ Unsure □

c. Other abnormal appearance:…………………………………………………………………………………..

6. Maloccluded incisors: Yes □ No □ Unsure □

a. Underbite/overbite: ……………………….

b. Estimation of size of malocclusion:……………mm

c. Overgrown incisors: Yes □ No □ Unsure □

i. If yes, estimated length of overgrowth:…………….mm

ii. Which incisors affected: Bottom □ Top □ Both□ Other □

d. Type of malocclusion: Early (bottom and top incisors meeting) □ Twisting malocclusion (incisors twisted) □ Separating incisors □

7. Elongated peg teeth: Yes □ No □ Unsure □

a. If yes, estimated length of overgrowth:……………mm

8. Variation in tooth height: Yes □ No □ Unsure □

a. If yes, how many teeth are of abnormal tooth height:…………..

9. Elongated molars: Yes □ No □ Unsure □

a. Estimation of length: ……..mm

10. Molar spurs: Yes □ No □ Unsure □

a. Number of teeth affected:……………

11. Molar sharpness: Yes □ No □ Unsure □

a. Number of teeth affected:……………

12. Tongue/cheek sores: Yes □ No □ Unsure □

a. If yes then current or scarred:……………….

b. If yes then how many are there?............

c. If current, how much of the tongue/cheek is affected:……………%

d. What depth do the sores affect: Deep □ Superficial □ Intermediate □ Unsure □

13. Fractured teeth: Yes □ No □ Unsure □

a. How many teeth are affected:…………….

14. Evidence of potential pain (struggling/refusal of exam): Yes □ No □ Unsure □

15. Other signs of dental disease:

a) Poor body condition (defined as BCS <2.5): Yes □ No □ Unsure □

b) Epiphora/ulceration below eye: Yes □ No □ Unsure □

c) Poor grooming (unkept coat): Yes □ No □ Unsure □

d) Hypersalivation/wet fur: Yes □ No □ Unsure □

e) Lack of coprophagia (faecal staining of perineum): Yes □ No □ Unsure □

f) Scattering of food around food bowl: Yes □ No □ Unsure (if pair housed) □

**Extra notes of interest on dental health**

…………………………………………………………………………………………………

…………………………………………………………………………………………………

…………………………………………………………………………………………………

…………………………………………………………………………………………………

…………………………………………………………………………………………………

…………………………………………………………………………………………………

…………………………………………………………………………………………………

……………………………………………………………

Part 4: Medical history

1. Previous dental disease diagnosed: Yes □ No □

a. If yes please state which:

…………………………………………………………………………………………………

…………………………………………………………………………………………..……

…..…………………………………………………………………………………..………

……………………………………………………………………………………………

b. If yes, what treatment given:

…………………………………………………………………………………………………

…………………………………………………………………………………………..……

…..……………………………………………………………………………………..………

………………………………………………………………………………………

c. Is the disease: Current & severe □ Current & mild □ Recurrent □ Resolved □ Unsure □

d. If no, was there a potential for dental disease due to clinical signs but it wasn't investigated?

i. Yes □ What clinical signs?.........................................................................................

2. Previous aural disease diagnosed: Yes □ No □

a. If yes please state which:

……………………………………………………………………………………………………

…………………………………………………………………………………………..……

…..……………………………………………………………………………………..………

………………………………………………………………………………………

b. If yes, what treatment given:

……………………………………………………………………………………………………

…………………………………………………………………………………………..……

…..……………………………………………………………………………………..………

………………………………………………………………………………………

c. Is the disease: Current & severe □ Current & mild □ Recurrent □ Resolved □ Unsure □

d. If no, was there a potential for aural disease due to clinical signs but it wasn't investigated?

i. Yes □ What clinical signs?.........................................................................................

**<u>Extra notes of interest on medical history</u>**

……………………………………………………………………………………………………

……………………………………………………………………………………………………

……………………………………………………………………………………………………
